# Automatic identification of small molecules that promote cell conversion and reprogramming

**DOI:** 10.1101/2020.04.01.021089

**Authors:** Francesco Napolitano, Trisevgeni Rapakoulia, Patrizia Annunziata, Akira Hasegawa, Melissa Cardon, Sara Napolitano, Lorenzo Vaccaro, Antonella Iuliano, Luca Giorgio Wanderlingh, Takeya Kasukawa, Diego L. Medina, Davide Cacchiarelli, Xin Gao, Diego di Bernardo, Erik Arner

## Abstract

Controlling cell fate has great potential for regenerative medicine, drug discovery, and basic research. Although numerous transcription factors have been discovered that are able to promote cell reprogramming and trans-differentiation, methods based on their up-regulation tend to show low efficiency. The identification of small molecules that can facilitate conversion between cell types can ameliorate this problem working through safe, rapid, and reversible mechanisms. Here we present DECCODE, an unbiased computational method for the identification of such molecules solely based on transcriptional data. DECCODE matches the largest available collection of drug-induced profiles (the LINCS database) for drug treatments against the largest publicly available dataset of primary cell transcriptional profiles (FANTOM5), to identify drugs that either alone or in combination enhance cell reprogramming and cell conversion. Extensive *in silico* and *in vitro* validation of DECCODE in the context of human induced pluripotent stem cells (hIPSCs) generation shows that the method is able to prioritize drugs enhancing cell reprogramming. We also generated predictions for cell conversion with single drugs and drug combinations for 145 different cell types and made them available for further studies.

## Introduction

Controlling cell fate has enormous potentials for regenerative medicine^1^, drug discovery^2^ and cell-based therapy^3^. A milestone discovery by Yamanaka and colleagues, who induced human stem cells via genetic reprogramming of mature somatic cells using four transcription factors (TFs)^4,5^, has recently revolutionized the field of stem cell biology. To date, numerous studies have revealed distinct sets of TFs that achieve or promote cell reprogramming^6^ and transdifferentiation^7,8^. However, these methods often suffer from low efficacy due to partly unknown barriers that need to be overcome for complete conversion^9^.

Optimizing the reprogramming system using non-invasive approaches such as small molecule treatment is a promising strategy that may increase the reprogramming potential. The cellular effects of small molecule treatment are often rapid, dose dependent and reversible^10^, and have potential for *in situ* regeneration therapeutic interventions^11^. Recently, several methods relying fully or partially on drug treatment to enhance cell conversion have emerged^12^. Many of these use fibroblasts as the starting cell type, reprogrammed towards pluripotency^13,14^ or trans-differentiated to specialized cell types including neurons^15^, endothelial cells^16^, pancreatic like cells^17^, cardiomyocytes^18^, hepatocytes^19^ or other cell types^20,21,22^. Such studies provide a proof of principle for drug-based reprogramming, the exact mechanisms of which, however, are often poorly understood, making extensive trial-and-error unavoidable. Indeed, methods for identifying small molecule candidates include either exhaustive screenings of drug libraries followed by marker gene readout^23^ or application of drugs known to modulate specific pathways involved in the desired lineage commitment^24^. While these methods are promising, they are laborious and do not scale.

Whereas computational approaches to identify novel combinations of transcription factors to facilitate cell reprogramming have been developed and validated^25,26^, no similar tools exist for small molecules. Here, we present a methodology to automatically identify small molecules that either alone or in combination enhance cell reprogramming and cell conversion. We analyzed 447 genome-wide expression profiles of untreated primary cells from the FANTOM5 project^27^ together with 107,404 transcriptional responses to small molecule treatment from the LINCS project^28^ to identify small molecules that drive the cell transcriptional program towards the one of the desired lineage. We make the results available in an online tool named DECCODE (<u>D</u>rug <u>E</u>nhanced <u>C</u>ell <u>CO</u>nversion using <u>D</u>ifferential <u>E</u>xpression), that when queried returns the top compounds predicted to enhance conversion towards the desired cell type. We extensively validated DECCODE to identify small molecules enhancing reprogramming of human fibroblasts towards human induced Pluripotent Stem Cells (hIPSCs). DECCODE is unbiased, as it does not rely on previous knowledge, and it can scale up to identify drugs to enhance cell conversion to any desired cell type. We make the results available in an online tool (available at the following URL: https://fantom.gsc.riken.jp/5/cellconv/) that when queried returns the top compounds predicted to enhance conversion towards a large collection of primary cell types.

## Results

### The DECCODE approach

A schematic representation of our approach is illustrated in **Figure 1A**. Given a target cell type, we constructed its cell type-specific differential gene expression profile using the FANTOM5 database^29^, the most extensive atlas of gene expression profiles across primary human cells^27^, thus obtaining 447 expression profiles corresponding to 145 different cell types (Methods). Specifically, the target cell-type expression profile was compared against the profiles of all the remaining cell-types to detect differentially expressed genes specific to the target cell-type. We then compared the target cell-type profile with drug-induced transcriptional profiles obtained from the LINCS database (GSE70138). LINCS contains 107,404 differential gene expression profiles corresponding to the transcriptional responses of 41 cell lines to 1,768 different drugs spanning different concentrations and time points, far exceeding any other publicly available resource of cellular perturbations^28^.

**Figure 1:**
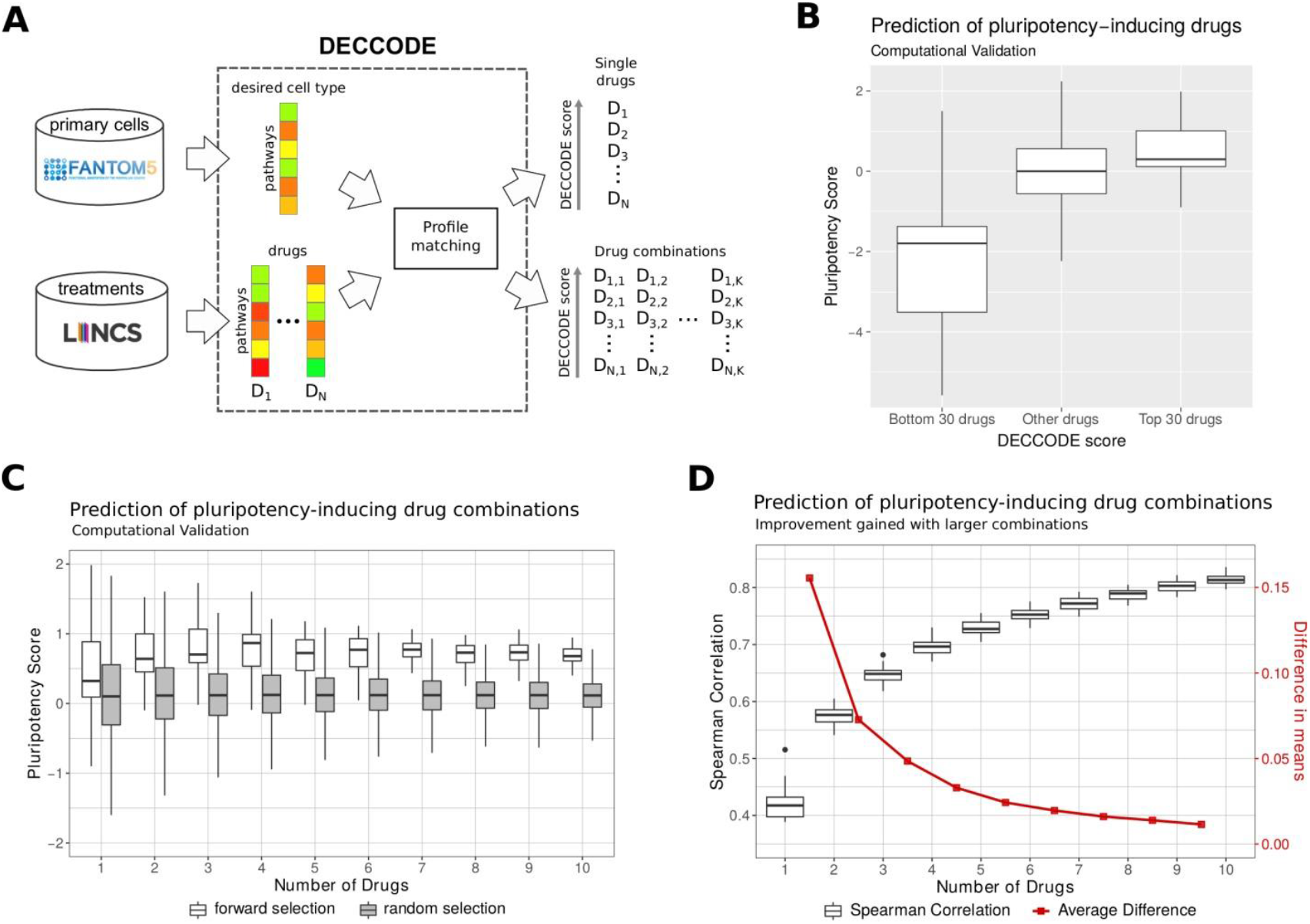
Computational identification of drugs facilitating cell conversion. (A) Workflow of the DECCODE approach. Target cell profiles are constructed from the FANTOM5 collection of human primary cell samples. Drug-induced consensus profiles are created for each of the treated cell lines included in the LINCS database. Single drugs or drug combinations are then prioritized based on their similarity with the target cell type profile. (B) In silico validation of single drugs facilitating conversion to hIPSCs. Drugs are grouped by their DECCODE scores and the Pluripotency Scores (PSs) of the drug-induced gene expression profiles within each group are computed and represented as a box-plot. (C) In silico validation of drug combinations of increasing size facilitating conversion to hIPSCs. PSs for drugs within each of the top 30 combinations are compared to random sets of the same size (see Methods for the details). Random selection was repeated 100 times. (D) Improvement obtained when using drug combinations of increasing size. The Spearman correlation between predicted drug combination profiles and hIPSC profile as more drugs are added is reported in the boxplot. The red line highlights the difference between the means of subsequent sets.

Since FANTOM5 and LINCS employ different expression profiling technologies, we first converted differential gene expression profiles (DGEPs) in both datasets to differential pathway-based expression profiles (DPEPs)^30^ (Methods) in order to enable an integrative analysis over the two datasets. Subsequently, we generated a consensus profile for each drug by merging together DPEPs across different time points and dosages. Finally, given a cell-type of interest, we searched among the 1,768 drugs that induce a transcriptional response similar to the expression profile of the target cell-type. The underlying hypothesis is that the selected drugs will induce a change in gene expression in the starting cell-type by making it more transcriptionally similar to the target cell-type, and thus facilitating the cell conversion process.

We also developed an extension of this method to predict drug combinations that synergize to enhance cell conversion. In previous work, we showed that combinatorial drug treatment is effectively described by a linear combination of the individual drug responses^31^ at the transcriptional level. The same finding has also been proven at the protein level, where protein dynamics in drug combinations can be explained by a linear superposition of their responses to individual drugs^32^. After confirming that the linear relationship also holds at the pathway level (Methods), we employed a multivariable linear regression model to describe the combined effect of drug combinations. First, for each drug, we selected the profile having highest DECCODE score across the treated cell lines, thus obtaining a single profile for each drug. Then we used forward selection to pick out the drug subsets yielding the most significant correlation with the target cell profile (Methods).

### Application of DECCODE to human IPS cells conversion

We first applied DECCODE in the single-drug mode to identify drugs enhancing cell reprogramming to human induced pluripotent stem cells (hIPSCs). We thus selected hIPSCs as the target cell-type and DECCODE returned the list of all 1,768 drugs ranked according to their predicted efficacy in enhancing cell reprogramming. We performed Drug Set Enrichment Analysis (DSEA)^30^ of the first 25 drugs in the ranking to identify those pathways that are consistently modulated by most of the drugs. As a result, we observed a consistent enrichment of pathways associated with pluripotency, such as differentiation and proliferation (**Supplementary Table 1**).

In order to further assess the validity of the DECCODE score, we devised an *in silico* validation method based on assigning a Pluripotency Score (PS) to each drug according to the upregulation of pluripotency specific genes and downregulation of somatic specific genes (Methods). We then compared the DECCODE scores with the PSs. A clear trend can be observed with top-ranked (higher DECCODE scores) drugs exhibiting higher PSs, and bottom-ranked (lower DECCODE scores) drugs exhibiting lower PSs, whereas no obvious correlation existed in the middle ranked profiles (**Figure 1B**). The full distribution of the DECCODE scores is reported in **Supplementary Figure 1**.

We then applied DECCODE in the drug-combination mode to identify drugs that can jointly enhance reprogramming to hIPSCs. Pluripotency Score and predicted similarity to the hIPSC profile of the top 30 drug combinations significantly improved when increasing the number of drugs, as assessed by Spearman correlation and adjusted r-squared. (**Figure 1C-D**, **Supplementary Figure 2A**). However, we observed that the most significant improvement was achieved when adding just one additional drug and gradually decreased as we kept adding more drugs, eventually reaching a plateau. Akaike Information Criterion (AIC) further confirmed that the relative goodness of fit increased more than what would be expected by chance as more drugs were added to the single drug models (**Supplementary Figure 2B**). The distribution of the transcriptional similarities between the two drug profiles in each of the top 30 drug-pair combinations was compared to randomly chosen drug-pairs (**Supplementary Figure 2C**). The results indicated that the two selected drugs in each combination tend to be transcriptionally different. Taken together, these *in silico* results suggest that drug combinations may offer an increased capacity to promote reprogramming compared to single drug administration.

### Experimental validation of DECCODE for conversion to hIPSCs

We set to experimentally validate predictions of DECCODE in single-drug mode applied to the hIPSCs reprogramming problem. As a biological model of reprogramming, we employed human secondary fibroblasts harboring a doxycycline-inducible OSKM (OCT4, SOX2, KLF4, C-MYC) gene cassette (hiF-T cells)^33^. We selected for experimental validation the top-ranked 25 drugs present in either of two widely used chemical screening libraries (Methods) that were predicted by DECCODE to enhance reprogramming to hIPSCs. In addition, we selected 20 drugs with scores in the lower half of the total ranking for comparison. hiF-T cells were treated with a total of 45 drugs in triplicate. After 21 days of treatment, cells were stained for the pluripotency marker TRA-1-60, imaged, and cell colony number and area were quantified for each well (see example in **Supplementary Figure 3**). The top-ranked drugs performed significantly better than the lower ranked drugs, either when considering the number of colonies or the total area covered by the colonies (**Figure 2A**). Several top-ranked drugs have been already associated with enhancement of reprogramming (**Supplementary Table 2**), including tranylcypromine, which we previously identified as a new positive regulator of the reprogramming process^33^.

**Figure 2:**
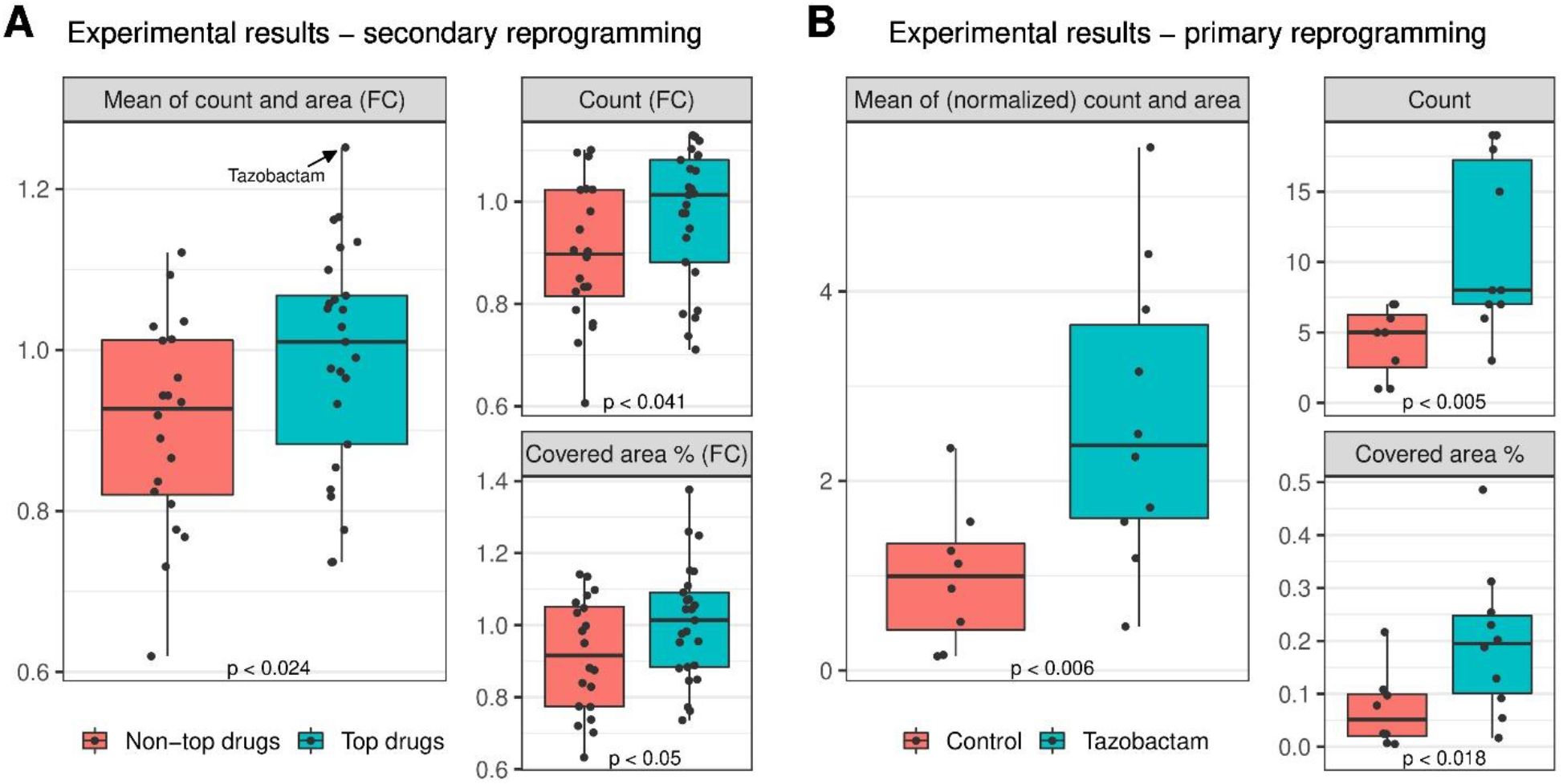
Experimental validation of DECCODE to identify drugs that enhance conversion to hIPSCs. (A) Secondary reprogramming: fold change (FC) relative to controls of the number of colonies and their total area following treatment either with the 25 drugs ranked by DECCODE at the top of the ranking (green boxes) or with 20 drugs ranked in the bottom half of the ranking (red boxes). Dots represent the effect of individual drugs in terms of the average fold change for three replicate experiments against controls. Main panel shows the combined average of both number of colonies (count) and their area expressed as FCs; smaller panels report counts and area separately. (B) Primary reprogramming: number of colonies and percentage of their total area following treatment with tazobactam and OSKM compared to OSKM alone (control). Dots represent the single values for each replicate.

Tazobactam, an antibiotic of the beta-lactamase inhibitor class previously unexamined in the context of cell reprogramming, achieved the highest performance when considering the area covered by the colonies and the second highest performance when considering the number of colonies (Figure 2A and **Supplementary Figure 4**), thus ranking first when considering both area and colony number together. Tazobactam was further validated by performing primary reprogramming of human primary foreskin fibroblasts through OSKM transduction either in the presence or absence of tazobactam. Both the number of colonies and the total area covered by the colonies confirmed the ability of tazobactam to enhance reprogramming to hIPSCs (**Figure 2B**).

### Resource

To create a comprehensive resource for drug-assisted cell conversions, we applied DECCODE to the whole FANTOM5 set of primary cells. We observed that closely related cell types exhibit high transcriptional similarity leading to tautological and nonspecific drug predictions. To reduce redundancy, we thus employed a two-level hybrid clustering of cell types taking into consideration both knowledge-driven and data-driven similarities (**Figure 3**). In the first level, we applied the Affinity Propagation^34^ algorithm to cluster primary cells using either an ontological similarity (Methods) or a transcriptional similarity, thus obtaining two different clusterings (**Figure 3**). For visualization purposes, we also performed a hierarchical clustering for both similarity measures (**Supplementary Figure 5, 6**). In the second level, cell types that were grouped together by both ontological and transcriptional clustering, were kept in one cluster, otherwise they were separated into distinct clusters. Finally, differential pathway-based expression profiles (DPEPs) of cell types in the same cluster were merged together to create a single consensus profile. We thus obtained 69 Consensus DPEPs corresponding to distinct “meta-cells”, i.e. an ensemble of different cell-types very similar in both ontological and transcriptional terms (the 69 meta-cell clusters are reported in **Supplementary Table 3**).

**Figure 3:**
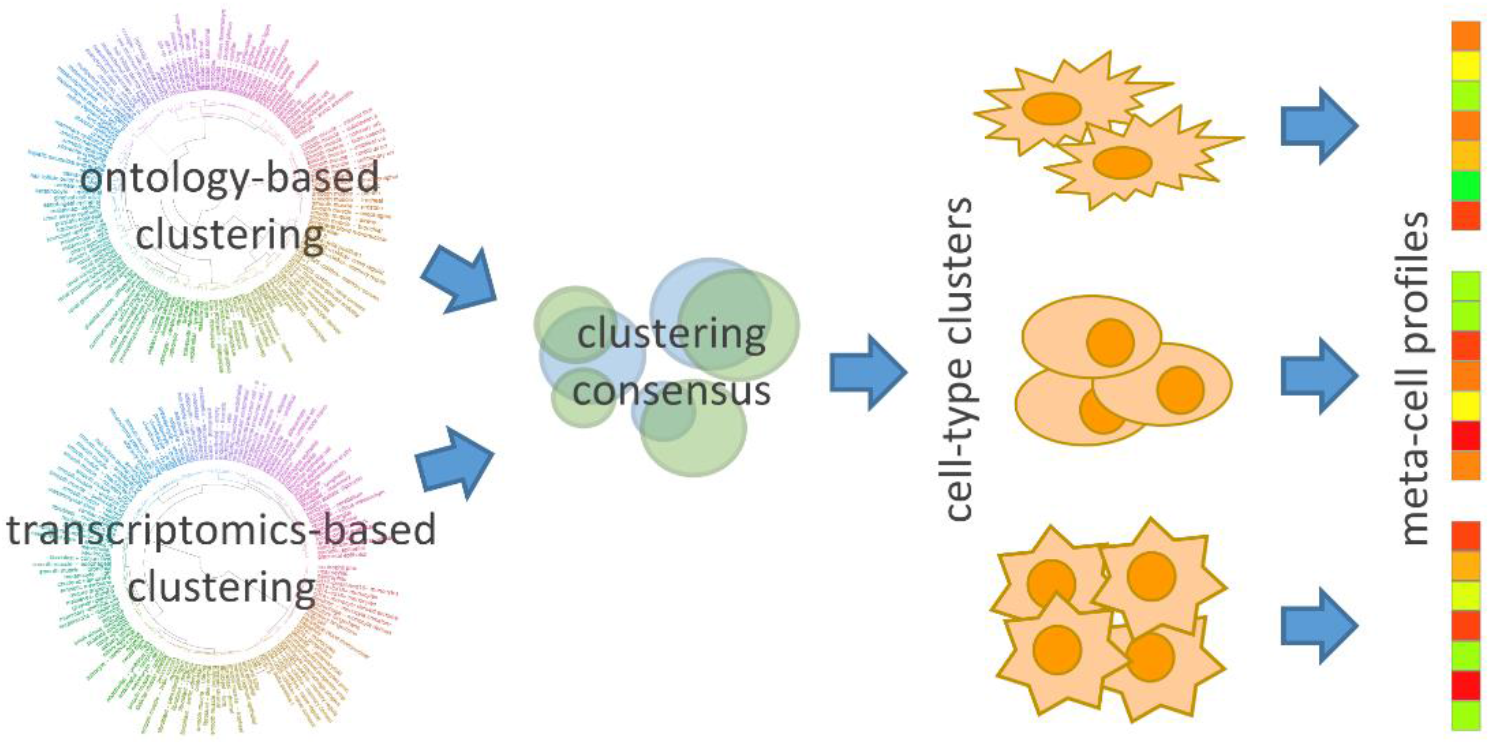
Two-level clustering to obtain cell-type specific consensus profiles.

We applied DECCODE to the 69 meta-cells profiles and found several drugs experimentally proved to facilitate cell conversions among the top 5% of ranked drugs (**Supplementary Tables 4&5**). Of note, two small molecules (Y-27632 and PD0325901) that were ranked among the top 20 candidates for conversion into the neuronal cell-type were previously experimentally proven to promote neural conversion. Y-27632, a ROCK inhibitor that assists in neuron survival, was used in combination with six other small molecules to convert human fibroblasts into neuronal-like cells^35^. PD0325901, a MEK-ERK inhibitor, facilitated the direct conversion of somatic cells to induced neuronal cells in a chemical cocktail of six compounds^36^.

Our method has been pre-computed on 69 meta-cells representing all the primary cell types included in the FANTOM5 database, and the top single- and multi-drug predictions are publicly available through the DECCODE web site (https://fantom.gsc.riken.jp/5/cellconv/) to provide an extensive resource that may support and complement future chemical enhanced cell conversion studies.

## Discussion

Therapies based on cell reprogramming and conversion are becoming a reality^37,38^ along with the need for methods that improve efficacy and safety of these processes. The use of small molecules, rather than genetic factors, is a promising approach to address safety issues. In addition to the ultimate goal of finding combinations that can fully convert one cell type to another, the identification of small molecules that can perform a partial conversion, or make the conversion more effective, also represents an important improvement on current methods. Here, we developed an unbiased method, DECCODE, which does not rely on expert knowledge of lineage-specific genes and pathways, scales to large numbers of cell conversions and drugs, and relies on publicly available data, thus not requiring massive screening efforts. Our method is the first validated computational approach for prioritizing small molecules promoting cell reprogramming. We have applied DECCODE to 145 human primary cell types from FANTOM5 and made the results available, providing a comprehensive resource including both single drugs and combinations of two drugs predicted to facilitate conversion to a variety of cell types. Together with such resources, we also released the full automated pipeline used for colony quantification, including source code and high resolution plate scans^39^ (DOI: 10.5281/zenodo.3732772).

Although the number of gene expression profiles following drug treatment available in public databases is substantial, the number of unique small molecules profiled is not. Indeed, only a subset of small molecules that have been experimentally validated to facilitate cell conversions, were transcriptionally profiled in LINCS. Moreover, considering that many of the profiled agents are kinase inhibitors, relevant for cancer therapy, publicly available drug profile resources represent a small portion of the “druggable genome” in cell conversion applications. Future profiling efforts that include additional libraries of small molecules will increase the utility of our approach.

Since our method relies on the analysis of transcriptomic data, there are some restrictions regarding the small molecules that can be captured. For example, epigenetic modifiers, such as HDAC inhibitors are extensively used in cell conversions to tackle the epigenetic barriers between different types of cells. The broad action and the nonspecific transcriptional behavior^40^ of these drugs limit their identifiability by DECCODE. In contrast, our approach gives priority to compounds exhibiting strong transcriptional regulation towards the target cell type. It may therefore be advisable to combine compounds identified in this study with treatment with epigenetic modifiers in order to further increase the efficacy of cell conversion.

Experiments on human fibroblasts confirmed the ability of DECCODE to predict small molecules facilitating cell reprogramming. The efficiency of reprogramming when treating cells with the best-ranked drugs was increased when compared to the control case. Our work demonstrates that DECCODE is able to distinguish and prioritize small molecules based on their potential to promote reprogramming and its usefulness in facilitating cell conversion should not be underestimated. We identified the core reprogramming chemicals for each lineage commitment, which could aid in establishing the role of various small molecules in different cell fates. We made our results available via a user-friendly interface to facilitate the design of cell conversion experiments involving chemical compounds. Our method provides a significant head start towards the development of systematic chemical based reprogramming strategies.

## Material and Methods

### Gene expression data

We used untreated primary cell profiles from the FANTOM5 collection and drug-induced gene expression profiles from the LINCS collection. We selected all primary human cells having at least two biological replicates from the FANTOM5 database (http://fantom.gsc.riken.jp/5/data), resulting in 447 samples, which correspond to 145 different cell types. Expression tables of robust CAGE peaks for these samples were processed as follows: we kept only the promoters located within 500 bp of known RefSeq transcripts (87,400 promoters). We added read counts of all the promoters sharing the same Entrez id annotation, resulting in 18,980 genes (**Supplementary Figure 7**). Read counts of samples were converted in CPM (counts per million) values and averaged across the same cell type. Z-score normalization was applied to each gene across cell types to obtain differential expression profiles for each cell type. Subsequently, genes in every cell type were ranked according to their expression, from the most expressed to the least expressed gene.

LINCS database is available as gene-based expression profiles. We downloaded the 5th level of differential gene expression signatures released on the GEO website (GEO accession GSE70138) which includes 107,404 profiles corresponding to 1,768 different drugs in 41 cell lines, 83 concentrations, and four treatment durations. Data access was performed through the cmapR package^41^. Genes in each drug profile were ranked according to their expression, from the most up-regulated to the most down-regulated gene.

### Conversion to pathway-based profiles

To harmonize the two datasets, we converted the ranked lists of genes from both cell-types (FANTOM5) and drug treatments (LINCS) into *pathway-based expression profiles* (PEPs). A PEP is a transcriptomic profile expressed in terms of pathways as opposed to genes. PEPs were introduced in our previous work^30^ and their efficacy for drug discovery applications was also proved^42^. To convert FANTOM5 and LINCS Gene Expression Profiles (GEPs) to PEPs, we applied the gep2pep Bioconductor package^43^ using all the 14,645 gene sets from 16 different gene set collections included in the MsigDB v6.1^44^. The gep2pep package iteratively performs Gene Set Enrichment Analysis (GSEA)^45^ to compute Enrichment Scores for each gene set and each expression profile. A PEP is then defined as a ranked list of pathways, each of which is associated with an Enrichment Score (and the corresponding p-value). Once FANTOM5 and LINCS GEPs are converted to PEPs, they can be directly compared (**Supplemental Figure 7**).

Various pathway-based profiles for the same gene expression profile can be obtained based on the chosen pathway database. In our case, as previously mentioned, we tried 16 different pathway collections available at the MSigDB database. We then evaluated which one out of these 16 collections best captured cell-type similarities, with respect to the Cell Ontology^46^. To this aim, we used the Cell Ontology annotation of cell-types created by the FANTOM5 consortium^47^. In order to obtain a numerical score for each pair of cell-types *i* and *j* in the ontology, we used the Jaccard Index as follows:

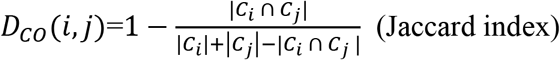

where C_i_ are the ontology ancestors of cell type *i*, *C*_*j*_ are the ontology ancestors of cell type *j*, *1* ≤ *i*, *j* ≤ *145*, *i* ≠ *j*. Then we defined the PEP-based distance between cell types *i* and *j* using the Manhattan distance as follows:

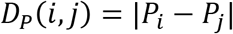

*P*_*i*_ is the PEP of cell type *i*, *P*_*j*_ is the PEP of cell type *j*, *1* ≤ *i*, *j* ≤ *145*, *i* ≠ *j*.

Finally, we compared the cell distances computed on the PEPs with the same cell distances based on the Cell Ontology (**Supplemental Figure 8**). The PEPs based on the C2 collection (Canonical Pathways) achieved the highest agreement with the ontology-based similarities, capturing more accurately the known cell hierarchy, when compared to a previously developed gene-based approach^48^.Thus, pathway-based profiles obtained with C2, which includes 250 pathways, were chosen for all further analyses.

### Merging of Pathway-based Expression Profiles

As previously proposed^48^, we merged multiple expression profiles elicited by the same drug treatment in order to obtain a single “consensus-profile” for each drug, thus enhancing drug-specific effects while reducing unrelated ones. The gep2pep package^43^ supports this operation by averaging the Enrichment Scores over multiple profiles and applying the Fisher method to aggregate their p-values. Using this approach, we merged together all the LINCS profiles induced by the same drug in the same cell line across different concentrations and treatment durations. An additional profile for each drug was generated by averaging all conditions, including different cell lines (termed “independent”). We used both approaches to obtain both cell-specific and cell-independent meta-profiles. We ended up with 17,259 drug-induced PEPs.

### Single-drug DECCODE scores

We converted LINCS and target cell-type PEPs to ranks based on their enrichment score (from the most enriched to the least enriched pathway). Then, we ranked each LINCS PEP by computing its L_1_ distance from the target cell-type PEP:

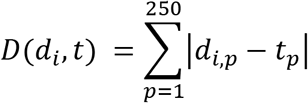

where *d_i_* is the LINCS PEP for drug *i*, *i* =1, …, 17,259 (number of drug-induced PEPs), *t* is the target cell type PEP, *p*=1,…, 250 (number of pathways in C2 collection). For further analysis, we considered only the top-ranked profile for each small molecule, resulting in 1,768 profiles (number of unique small molecules).

### In silico validation with the Pluripotency Score

While the DECCODE framework is based on an unbiased, data-driven approach, we devised a pluripotency-specific method to score gene expression profiles based on prior knowledge about genes involved in the conversion to hIPSCs. We then compared these scores with DECCODE scores to validate the predictions. The pluripotency score (PS) is based on genes that were identified as differentially expressed during reprogramming. In particular, we used the “early pluripotency”, “late pluripotency”, “early somatic”, and “late somatic” gene sets previously identified^33^ that characterize gene expression dynamics in the corresponding stages of conversion from human fibroblasts (HiF-T) to hIPSCs. The original study included also other six sets from the same context, which we used as statistical background. For each of the ten sets and for each drug-induced gene expression profile, we computed an Enrichment Score (ES) using the gep2pep tool. We then ranked them from 1^st^ to 10^th^ according to their ESs, thus obtaining a PEP profile. Finally, we computed the Pluripotency Score (PS) for each profile *p* as:

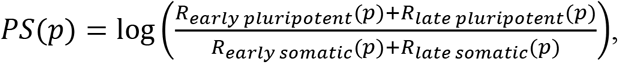

where R_x_(p) is the rank of the gene set *x* within the profile *p*. The score is positive (negative) when genes associated with pluripotency stages tend to be more up-regulated (down-regulated) than genes associated with somatic stages.

### DECCODE scores for drug combinations

We showed in previous work that the transcriptional response to combinatorial drug treatment at promoters and enhancers is effectively described by a linear combination of the responses of the individual drugs (log2FC values)^31^. We used our previous dataset to test if this additive relationship also applies to PEPs. Accordingly, we performed multivariable linear regression analysis, where PEPs of individual drugs were considered as explanatory variables and the PEP of combinatorial drug action as the response variable. We applied our analysis to five pathway databases, Biological Process (BP), Molecular Function (MF), Cellular Component (CC), Transcription Factor Targets (TFT) and Canonical Pathways (C2_CP). **Supplementary Table 6** demonstrates the performance of the linear regression model after ten-fold cross-validation in all the three drug combinations and the four pathway collections. The results show that the linear model using PEPs can describe the relation between single and combinatorial treatment.

To validate whether both single drug PEPs contribute to the model, we performed the same regression analysis 100,000 times with random permutations of one of the single drug PEP. The Pearson correlation between the observed and predicted values after the permutations was significantly lower for all combinations compared to the regression model based on the non-permuted individual drug PEPs (**Supplementary Figure 9**).

In order to produce DECCODE scores for drug combinations, we considered only the top-ranked PEP for each drug based on its Manhattan distance to the target cell type. For each drug PEP (1,768), we fitted a simple linear regression model and we ranked drugs based on the spearman correlation between fitted and observed values. We picked the top 30 drug PEPs and searched through the remaining drugs to find out which one should be added to the current models to best improve the Spearman correlation. Repeated occurrences of the same drug sets in different order were discarded. We continued to add variables to the top 30 models until we reached 10 predictors. Computational validation of the obtained combinations was assessed using PSs (Figure 1B). In particular, for any drug combination the median of the corresponding PSs was used. Moreover, the top 30 solutions were considered for a given drug combination size, thus obtaining 30 median PS values. In order to obtain a corresponding null distribution, the same calculation was performed also for random drug combinations of equal size. The random selection was repeated 100 times for each size, thus obtaining 100 times 30 median PSs. Figure 1B summarizes this analysis by reporting the obtained 30 versus 300 median PSs for drug combinations of size 1 to 10.

### Selection of the drugs for the experimental validation

In order to validate the method experimentally, we selected 25 drugs from the top of the DECCODE ranking, plus 20 non-top drugs for comparison. In particular, to build the set of non-top drugs, we chose 10 drugs from the middle of the ranking and 10 drugs from the bottom. In case of an overlap between top and non-top drugs due to the same drugs being profiled across multiple cellular contexts in the LINCS database, we removed the repeated drugs from the non-top sets and chose the next one in the ranking. In all cases, only the drugs included in the Prestwick-FDA library or in the SelleckChem Kinase inhibitors library were considered.

### Human Cellular Reprogramming

All the reprogramming experiments and procedures were performed as previously described^33^. In summary, secondary reprogramming was performed by seeding, on a confluent irradiated MEF-feeder layer, clonal TERT-immortalized secondary fibroblasts harboring a doxycycline-inducible OSKM cassette. The day after seeding reprogramming was initiated by doxycycline supplementation and protracted for 21 days.

For primary reprogramming, BJ foreskin fibroblasts were infected with a lentivirus harboring the constitutive OSKM cassette (pLM-fSV2A)^49^ split onto an irradiated MEF-feeder layer and reprogrammed for 15 days.

At the end of each reprogramming, quantitative analysis of colony number and area was performed using a TRA-1-60 chromogenic staining in bright field. All candidate drugs for reprogramming were tested for the entire duration of the reprogramming process at a final concentration of 10nM in several technical or biological replicates, as indicated.

### Colony quantification

To quantify colony number and size in an unbiased and reproducible way, a completely automated procedure was developed, which is divided in two phases. The first phase was performed through a Matlab script which identifies each well inside all the plate scans, applies a 3X contrast, and saves each of them to a separate image file. The second phase was performed by an ImageJ macro that loads the well images produced by the previous step and performs the final counting and area estimation on each of them. Both Matlab and ImageJ source code, together with the high resolution plate scan images, are available online^39^ (DOI: 10.5281/zenodo.3732772).

In secondary reprogramming experiments, colony count and area values were averaged across the three replicates of the same treatment and across the two controls on the same plate. Average fold change of treatments versus controls were then obtained accordingly (**Figure 2A**, small panels).

In order to summarize both count and covered area values together, the corresponding fold changes were averaged (**Figure 2A**, main panel). In the primary reprogramming experiment, counts and areas for tazobactam treatment and controls were compared directly (**Figure 2B**, small panels). Two controls were excluded according to the Bonferroni Outlier Test (*p* < 0.0118 and *p* < 0.0106 respectively). In the case of primary reprogramming results, in order to summarize counts and covered area together, all the absolute values were normalized dividing by the corresponding mean of the controls (**Figure 2B**, main panel).

### Computation of DECCODE scores for all the FANTOM5 cell types

The FANTOM5 cell types include sub-types that are very similar, thus the corresponding expression profile are not different enough to produce sub-type specific predictions. Therefore, we merged similar cell types to form a single meta-cell profile (see methods subsection “Merging of Pathway-based Expression Profiles”). In order to systematically select which cell-type profiles to merge, we took advantage of the previously computed PEP-based and ontology-based cell type distances (refer to subsection “Conversion to pathway-based profiles”). We applied the Affinity Propagation algorithm^34^ individually to each of the two pairwise distances to obtain two different clusterings of the same cell types (**Supplementary Figures 5-6**). Affinity Propagation clustering was performed using the “apcluster” R package^50^. Finally, we built a consensus clustering by assigning two cell types to the same cluster if and only if they were assigned to the same cluster by both the ontology-based and PEP-based clusterings. Meta-cell profiles are obtained by merging all the profiles included in the same cluster. We then computed single-drug and multiple-drug DECCODE scores for all the meta-cell profiles.

## Acknowledgements

EA was supported by a Research Grant from MEXT to the RIKEN Center for Integrative Medical Sciences. XG was supported by funding from King Abdullah University of Science and Technology (KAUST), under award number FCC/1/1976-18-01, FCC/1/1976-23-01, FCC/1/1976-25-01, FCC/1/1976-26-01, and FCS/1/4102-02-01. DLM was supported by the Italian Telethon Foundation under the project number TMDMCBX16TT. DC was supported by Fondazione Telethon Core Grant, Armenise-Harvard Foundation Career Development Award, European Research Council (grant agreement 759154, CellKarma), and the Rita-Levi Montalcini program from MIUR.

## Author contributions

Data analysis was performed by FN and TR and supervised by XG, DdB and EA. Additional analysis was performed by SN and AI. Validation experiments were performed by PA, LV and LGW, supervised by DC and DLM. The database and website were implemented by MC and AH, and supervised by TK. The manuscript was drafted by TR, FN, DdB and EA with additional input from all other co-authors. The study was conceived by EA and jointly supervised by EA, DdB, DC and XG.

## References

1. Cohen, D. E. & Melton, D. Turning straw into gold: directing cell fate for regenerative medicine. Nat. Rev. Genet. 12, 243–52 (2011).

2. Avior, Y., Sagi, I. & Benvenisty, N. Pluripotent stem cells in disease modelling and drug discovery. Nat. Rev. Mol. Cell Biol. 17, 170–182 (2016).

3. Kikuchi, T. et al. Human iPS cell-derived dopaminergic neurons function in a primate Parkinson’s disease model. Nature 548, 592–596 (2017).

4. Takahashi, K. & Yamanaka, S. Induction of Pluripotent Stem Cells from Mouse Embryonic and Adult Fibroblast Cultures by Defined Factors. Cell 126, 663–676 (2006).

5. Takahashi, K. et al. Induction of Pluripotent Stem Cells from Adult Human Fibroblasts by Defined Factors. Cell 131, 861–872 (2007).

6. Soufi, A. et al. Pioneer Transcription Factors Target Partial DNA Motifs on Nucleosomes to Initiate Reprogramming. Cell 161, 555–568 (2015).

7. Rosa, F. F. et al. Direct reprogramming of fibroblasts into antigen-presenting dendritic cells. Sci. Immunol. 3, eaau4292 (2018).

8. Sekiya, S. & Suzuki, A. Direct conversion of mouse fibroblasts to hepatocyte-like cells by defined factors. Nature 475, 390–393 (2011).

9. Smith, Z. D., Sindhu, C. & Meissner, A. Molecular features of cellular reprogramming and development. Nat. Rev. Mol. Cell Biol. 17, 139–54 (2016).

10. Zhang, Y., Li, W., Laurent, T. & Ding, S. Small molecules, big roles -- the chemical manipulation of stem cell fate and somatic cell reprogramming. J. Cell Sci. 125, 5609–20 (2012).

11. Biswas, D. & Jiang, P. Chemically Induced Reprogramming of Somatic Cells to Pluripotent Stem Cells and Neural Cells. Int. J. Mol. Sci. 17, 226 (2016).

12. Federation, A. J., Bradner, J. E. & Meissner, A. The use of small molecules in somatic-cell reprogramming. Trends Cell Biol. 24, 179–87 (2014).

13. Zhu, S. et al. Reprogramming of Human Primary Somatic Cells by OCT4 and Chemical Compounds. Cell Stem Cell 7, 651–655 (2010).

14. Hou, P. et al. Pluripotent Stem Cells Induced from Mouse Somatic Cells by Small-Molecule Compounds. Science (80-.). 341, 651–654 (2013).

15. Ladewig, J. et al. Small molecules enable highly efficient neuronal conversion of human fibroblasts. Nat. Methods 9, 575–578 (2012).

16. Sayed, N. et al. Transdifferentiation of Human Fibroblasts to Endothelial Cells: Role of Innate Immunity. Circulation 131, 300–309 (2015).

17. Zhu, S. et al. Human pancreatic beta-like cells converted from fibroblasts. Nat. Commun. 7, 10080 (2016).

18. Cao, N. et al. Conversion of human fibroblasts into functional cardiomyocytes by small molecules. Science (80-.). 352, 1216–1220 (2016).

19. Lim, K. T. et al. Small Molecules Facilitate Single Factor-Mediated Hepatic Reprogramming. Cell Rep. 15, 814–829 (2016).

20. Li, J. et al. Artemisinins Target GABA A Receptor Signaling and Impair α Cell Identity. Cell 168, 86–100.e15 (2017).

21. Wang, Y. et al. Conversion of Human Gastric Epithelial Cells to Multipotent Endodermal Progenitors using Defined Small Molecules. Cell Stem Cell 19, 449–461 (2016).

22. Cheng, L. et al. Direct conversion of astrocytes into neuronal cells by drug cocktail. Cell Res. 25, 1269–1272 (2015).

23. Li, W., Li, K., Wei, W. & Ding, S. Chemical approaches to stem cell biology and therapeutics. Cell Stem Cell 13, 270–83 (2013).

24. Federation, A. J., Bradner, J. E. & Meissner, A. The use of small molecules in somatic-cell reprogramming. Trends Cell Biol. 24, 179–187 (2014).

25. Cahan, P. et al. CellNet: Network Biology Applied to Stem Cell Engineering. Cell 158, 903–915 (2014).

26. Rackham, O. J. L. et al. A predictive computational framework for direct reprogramming between human cell types. Nat. Genet. 48, 331–335 (2016).

27. Forrest, A. R. R. et al. A promoter-level mammalian expression atlas. Nature 507, 462–70 (2014).

28. Keenan, A. B. et al. The Library of Integrated Network-Based Cellular Signatures NIH Program: System-Level Cataloging of Human Cells Response to Perturbations. Cell Syst. 6, 13–24 (2018).

29. Noguchi, S. et al. FANTOM5 CAGE profiles of human and mouse samples. Sci. Data 4, 170112 (2017).

30. Napolitano, F., Sirci, F., Carrella, D. & Di Bernardo, D. Drug-set enrichment analysis: A novel tool to investigate drug mode of action. Bioinformatics 32, 235–241 (2016).

31. Rapakoulia, T. et al. Genome-scale regression analysis reveals a linear relationship for promoters and enhancers after combinatorial drug treatment. Bioinformatics 33, (2017).

32. Geva-Zatorsky, N. et al. Protein dynamics in drug combinations: a linear superposition of individual-drug responses. Cell 140, 643–51 (2010).

33. Cacchiarelli, D. et al. Integrative analyses of human reprogramming reveal dynamic nature of induced pluripotency. Cell 162, 412 (2015).

34. Frey, B. J. & Dueck, D. Clustering by Passing Messages Between Data Points. Science (80-.). 315, 972–976 (2007).

35. Hu, W. et al. Direct Conversion of Normal and Alzheimer’s Disease Human Fibroblasts into Neuronal Cells by Small Molecules. Cell Stem Cell 17, (2015).

36. Dai, P., Harada, Y. & Takamatsu, T. Highly efficient direct conversion of human fibroblasts to neuronal cells by chemical compounds. J. Clin. Biochem. Nutr. 56, 166–170 (2015).

37. Stoddard-Bennett, T. & Reijo Pera, R. Treatment of Parkinson’s Disease through Personalized Medicine and Induced Pluripotent Stem Cells. Cells 8, (2019).

38. Mandai, M. et al. Autologous Induced Stem-Cell–Derived Retinal Cells for Macular Degeneration. N. Engl. J. Med. 376, 1038–1046 (2017).

39. Napolitano, F. et al. Automatic identification of small molecules that promote cell conversion and reprogramming - plate scans, colony quantification scripts, and DECCODE ranking. (2020). doi:10.5281/ZENODO.3732772

40. Chen, H. P., Zhao, Y. T. & Zhao, T. C. Histone deacetylases and mechanisms of regulation of gene expression. Crit. Rev. Oncog. 20, 35–47 (2015).

41. Enache, O. M. et al. The GCTx format and cmap{Py, R, M, J} packages: resources for optimized storage and integrated traversal of annotated dense matrices. Bioinformatics 35, 1427–1429 (2019).

42. Napolitano, F. et al. gene2drug: a computational tool for pathway-based rational drug repositioning. Bioinformatics 34, 1498–1505 (2018).

43. Napolitano, F., Carrella, D., Gao, X. & di Bernardo, D. gep2pep: a bioconductor package for the creation and analysis of pathway-based expression profiles. Bioinformatics (2019). doi:10.1093/bioinformatics/btz803

44. Liberzon, A. et al. The Molecular Signatures Database (MSigDB) hallmark gene set collection. Cell Syst. 1, 417–425 (2015).

45. Subramanian, A. et al. Gene set enrichment analysis: A knowledge-based approach for interpreting genome-wide expression profiles. Proc. Natl. Acad. Sci. 102, 15545–15550 (2005).

46. Bard, J., Rhee, S. Y. & Ashburner, M. An ontology for cell types. Genome Biol. 6, (2005).

47. Lizio, M. et al. Gateways to the FANTOM5 promoter level mammalian expression atlas. Genome Biol. 16, 22 (2015).

48. Iorio, F. et al. Discovery of drug mode of action and drug repositioning from transcriptional responses. Proc. Natl. Acad. Sci. U. S. A. 107, 14621–6 (2010).

49. Papapetrou, E. P. et al. Genomic safe harbors permit high β-globin transgene expression in thalassemia induced pluripotent stem cells. Nat. Biotechnol. 29, 73–81 (2011).

50. Bodenhofer, U., Kothmeier, A. & Hochreiter, S. APCluster: an R package for affinity propagation clustering. Bioinformatics 27, 2463–2464 (2011).

