## Supplemental Material for "Automatic identification of small molecules that promote cell conversion and reprogramming"

### Supplementary Materials

**Supplementary Table 1:** Results of the Drug Set Enrichment Analysis[1] applied to the set of top drugs (25 drugs with highest DECCODE scores). The table reports the top 30 pathways identified in the Gene Ontology – Biological Process collection out of 4,435 included in MSigDB v6.1. According to the DSEA, such pathways represent direct or indirect targets of most drugs in the top 25 set. The Enrichment Scores (ES) and the corresponding p-values respectively report the magnitude and the significance of the association. All top 30 pathways appear down-regulated (negative ES) and many of them are involved in cell proliferation and differentiation, which is coherent with the predicted effect of pluripotency induction.

| Rank | GO Term | Description | ES | p-value |
|---|---|---|---|---|
| 1 | Regulation of cytokine biosynthetic process | Any process that modulates the frequency, rate or extent of the chemical reactions and pathways resulting in the formation of cytokines. | -0.816 | $7.1054^{-15}$ |
| 2 | Myeloid leukocyte differentiation | The process in which a relatively unspecialized myeloid precursor cell acquires the specialized features of any cell of the myeloid leukocyte lineage. | -0.795 | $4.05231^{-14}$ |
| 3 | Regulation of cell activation | Any process that modulates the frequency, rate or extent of cell activation, the change in the morphology or behavior of a cell resulting from exposure to an activating factor such as a cellular or soluble ligand. | -0.736 | $3.64975^{-12}$ |
| 4 | Response to molecule of bacterial origin | Any process that results in a change in state or activity of an organism (in terms of movement, secretion, enzyme production, gene expression, etc.) as a result of a stimulus by molecules of bacterial origin such as peptides derived from bacterial flagellin. | -0.732 | $4.72367^{-12}$ |
| 5 | Leukocyte differentiation | The process in which a relatively unspecialized hemopoietic precursor cell acquires the specialized features of a leukocyte. A leukocyte is an achromatic cell of the myeloid or lymphoid lineages capable of ameboid movement, found in blood or other tissue. | -0.725 | $8.02824^{-12}$ |
| 6 | Osteoclast differentiation | The process in which a relatively unspecialized monocyte acquires the specialized features of an osteoclast. An osteoclast is a specialized phagocytic cell associated with the absorption and removal of the mineralized matrix of bone tissue. | -0.720 | $1.14407^{-11}$ |
| 7 | Regulation of cytokine production | Any process that modulates the frequency, rate, or extent of production of a cytokine. | -0.716 | $1.56148^{-11}$ |
| 8 | Inflammatory response | The immediate defensive reaction (by vertebrate tissue) to infection or injury caused by chemical or physical agents. The process is characterized by local vasodilation, extravasation of plasma into intercellular spaces and accumulation of white blood cells and macrophages. | -0.713 | $1.87594^{-11}$ |
| 9 | Positive regulation of cell activation | Any process that activates or increases the frequency, rate or extent of activation. | -0.713 | $1.95929^{-11}$ |
| 10 | Leukocyte migration | The movement of a leukocyte within or between different tissues and organs of the body. | -0.708 | $2.77252^{-11}$ |
| 11 | Regulation of b cell proliferation | Any process that modulates the frequency, rate or extent of B cell proliferation. | -0.707 | $2.88425^{-11}$ |
| 12 | Regulation of leukocyte proliferation | Any process that modulates the frequency, rate or extent of leukocyte proliferation. | -0.707 | $2.96568^{-11}$ |
| 13 | Positive regulation of cytokine biosynthetic process | Any process that activates or increases the frequency, rate or extent of the chemical reactions and pathways resulting in the formation of cytokines. | -0.706 | $3.07102^{-11}$ |

| | | | | |
|---|---|---|---|---|
| 14 | Adaptive immune response | An immune response based on directed amplification of specific receptors for antigen produced through a somatic diversification process, and allowing for enhanced response to subsequent exposures to the same antigen (immunological memory). | -0.705 | $3.36277^{-11}$ |
| 15 | Regulation of leukocyte differentiation | Any process that modulates the frequency, rate or extent of leukocyte differentiation. | -0.696 | $6.36902^{-11}$ |
| 16 | Myeloid leukocyte activation | A change in the morphology or behavior of a myeloid leukocyte resulting from exposure to an activating factor such as a cellular or soluble ligand. | -0.694 | $7.40514^{-11}$ |
| 17 | Regulation of peptidyl tyrosine phosphorylation | Any process that modulates the frequency, rate or extent of the phosphorylation of peptidyl-tyrosine. | -0.690 | $9.29625^{-11}$ |
| 18 | Positive regulation of cytokine production | Any process that activates or increases the frequency, rate or extent of production of a cytokine. | -0.689 | $1.03853^{-10}$ |
| 19 | Regulation of blood vessel endothelial cell migration | Any process that modulates the frequency, rate or extent of the migration of the endothelial cells of blood vessels. | -0.688 | $1.07958^{-10}$ |
| 20 | Positive regulation of cell cell adhesion | Any process that activates or increases the rate or extent of cell adhesion to another cell. | -0.686 | $1.23886^{-10}$ |
| 21 | Cell activation | A change in the morphology or behavior of a cell resulting from exposure to an activating factor such as a cellular or soluble ligand. | -0.686 | $1.25275^{-10}$ |
| 22 | Regulation of cell cell adhesion | Any process that modulates the frequency, rate or extent of attachment of a cell to another cell. | -0.681 | $1.7974^{-10}$ |
| 23 | Positive regulation of blood vessel endothelial cell migration | Any process that activates or increases the frequency, rate or extent of the migration of the endothelial cells of blood vessels. | -0.679 | $2.01367^{-10}$ |
| 24 | Regulation of interferon gamma production | Any process that modulates the frequency, rate, or extent of interferon-gamma production. Interferon-gamma is also known as type II interferon. | -0.679 | $2.04494^{-10}$ |
| 25 | Response to bacterium | Any process that results in a change in state or activity of a cell or an organism (in terms of movement, secretion, enzyme production, gene expression, etc.) as a result of a stimulus from a bacterium. | -0.677 | $2.28166^{-10}$ |
| 26 | Positive regulation of cell adhesion | Any process that activates or increases the frequency, rate or extent of cell adhesion. | -0.677 | $2.33346^{-10}$ |
| 27 | Regulation of homotypic cell cell adhesion | Any process that modulates the frequency, rate, or extent of homotypic cell-cell adhesion. | -0.670 | $3.75381^{-10}$ |
| 28 | Innate immune response | Innate immune responses are defense responses mediated by germline encoded components that directly recognize components of potential pathogens. | -0.669 | $4.05845^{-10}$ |
| 29 | Positive regulation of peptidyl tyrosine phosphorylation | Any process that activates or increases the frequency, rate or extent of the phosphorylation of peptidyl-tyrosine. | -0.668 | $4.23039^{-10}$ |
| 30 | Negative regulation of b cell proliferation | Any process that stops, prevents or reduces the rate or extent of B cell proliferation. | -0.664 | $5.38032^{-10}$ |

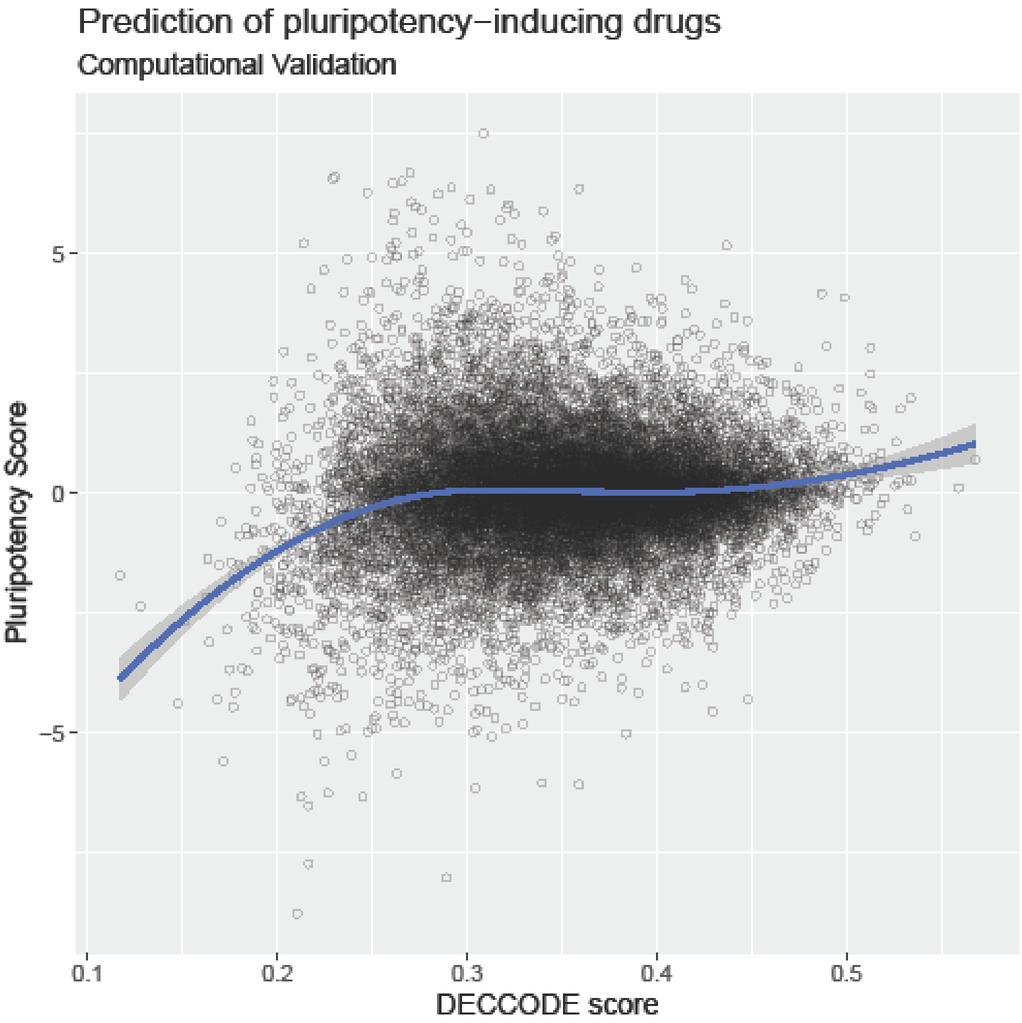


**Supplementary Figure 1:** DECCODE scores are evaluated against the PSs of drugs. Top-ranked (higher DECCODE scores) drugs exhibit higher PSs while bottom-ranked drugs exhibit lower PSs.

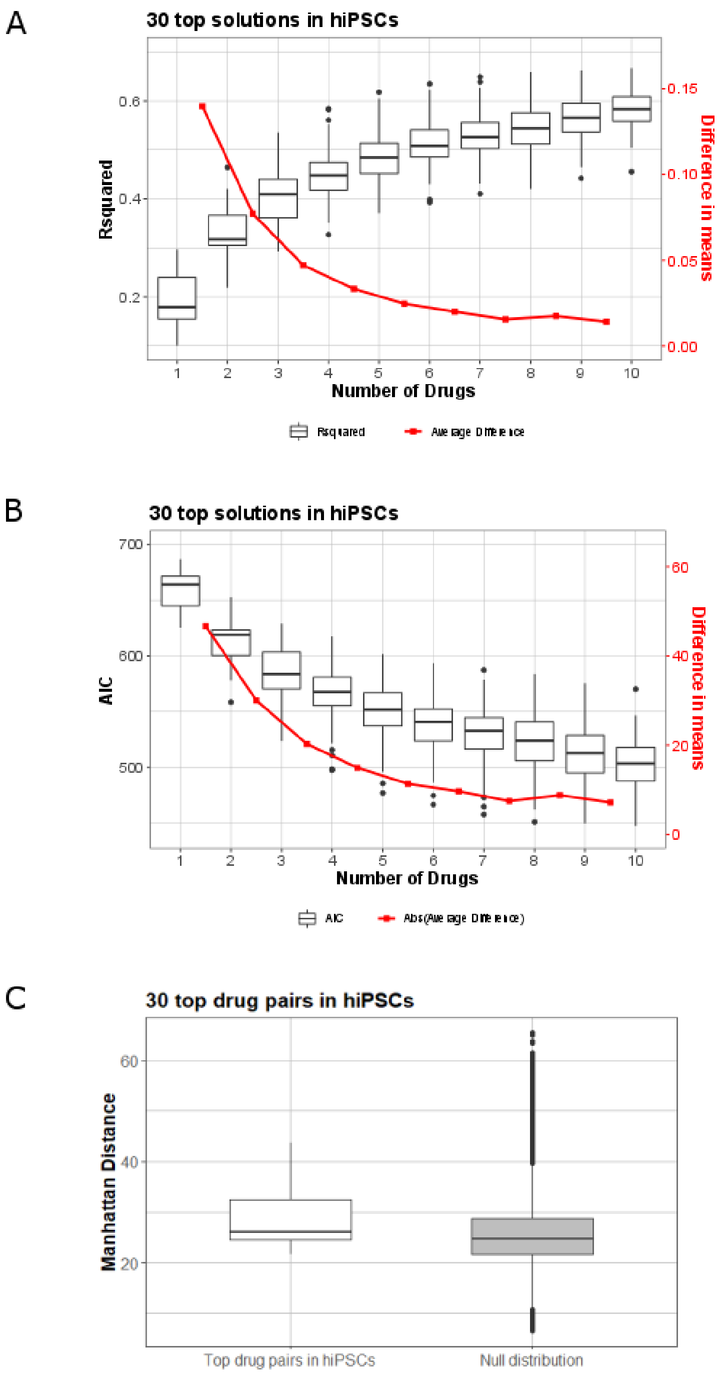


**Supplementary Figure 2**: A) Rsquared and B) AIC criterion of the top 30 regression solutions for the hIPSC target profile as more drugs are added to the regression models. Red line highlights the average incremental improvement. C) Distribution of the distances between the drug profiles for the top 30 selected drug pairs in hiPSCs by DECCODE. Null distribution was created by random sampling 1000 drug profiles from LINCS dataset and computing their pairwise distances (499500 distances).

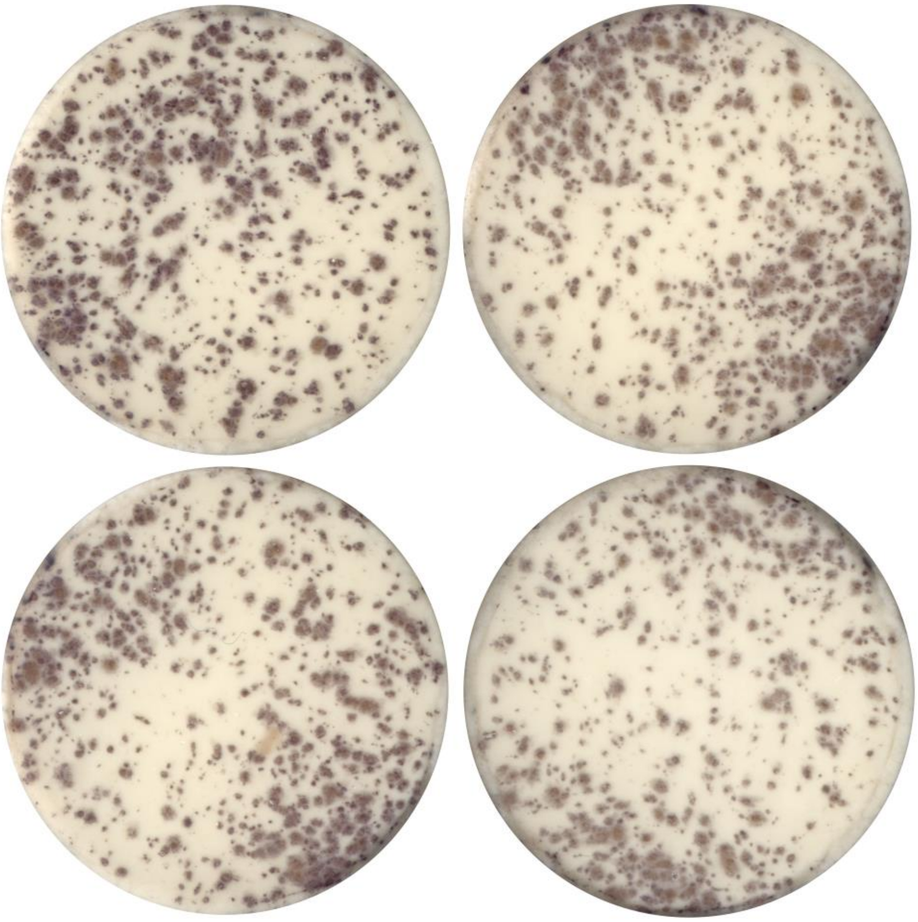

**Supplementary Figure 3**: Imaging of wells treated with tazobactam and OSKM (left) against the controls (only OSKM) for the specific plate (right). The fold change in number of colonies and area covered by the colonies for each drug treatment was computed against the control experiments of the corresponding plate.

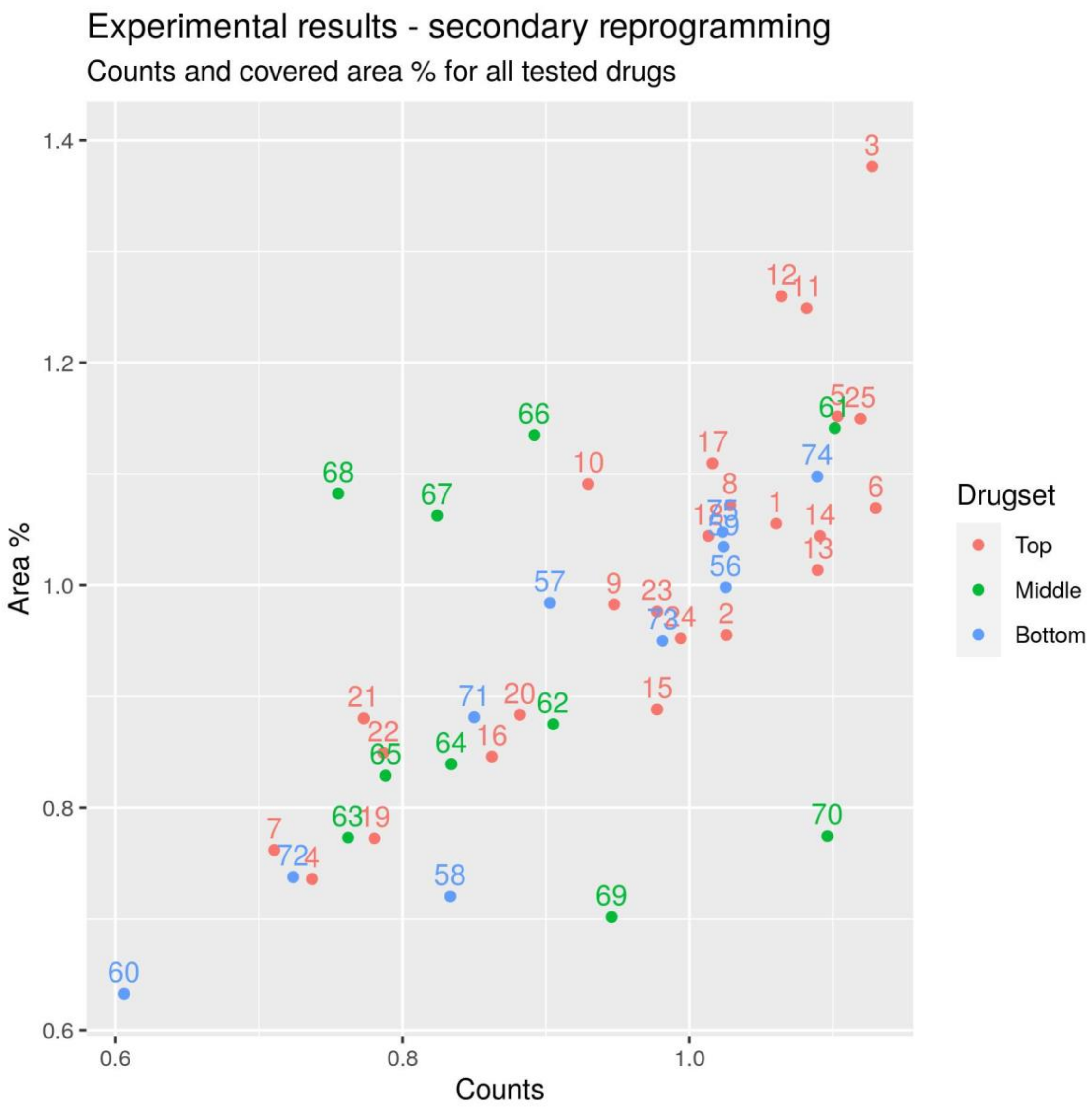


**Supplementary Figure 4**: Experimental validation of drugs enhancing conversion to hIPSCs: number of colonies formed versus % of covered area. Tazobactam (ID: 3) shows the highest performance for covered

**Supplementary Table 2:** Top-ranked drugs by DECCODE for the hIPSCs target profile. Several drugs have been already associated with enhancement of the reprogramming process.

| Drugs | Indication | ID |
|---|---|---|
| **Motesanib**[2] | treatment in solid tumors | 1 |
| **Fluticasone** | activating glucocorticoid receptors, inhibiting nuclear factor kappa b and inhibiting lung eosinophilia in rats | 2 |
| **Tazobactam** | bacterial β-lactamase inhibitor | 3 |
| **Cyclizine** | histamine H1 antagonist | 4 |
| **Etofenamate**[3] | nonsteroidal anti-inflammatory drug (NSAID), COX inhibitor | 5 |
| **Pentoxifylline** | modulates immunologic activity by stimulating cytokine production. | 6 |
| **Irsogladine** | anti-inflammatory agent | 7 |
| **Leflunomide** | pyrimidine synthesis inhibitor/chemotherapeutic | 8 |
| **Dexfenfluramine** | serotonergic anorectic drug/studied in obesity | 9 |
| **Paroxetine** | selective serotonin reuptake inhibitor (SSRI) drug commonly known as Paxil | 10 |
| **Afatinib** | tyrosine kinase inhibitor /ErbB family blocker | 11 |
| **Doramapimod** | highly potent p38 MAPK inhibitor | 12 |
| **Nalbuphine** | anticonvulsant effect/inhibited breast cancer cell growth and tumorigenesis | 13 |
| **PIK-93** | PI4KIIIβ inhibitor | 14 |
| **Glycopyrrolate** | synthetic anticholinergic agent | 15 |
| **SGX523** | MET receptor tyrosine kinase inhibitor. | 16 |
| **Dasatinib**[4] | Src family tyrosine kinase inhibitor | 17 |
| **SB-203580**[5] | inhibitor of p38α and p38β | 18 |
| **Doxycycline**[6] | antibacterial agent | 19 |
| **Saracatinib**[7] | inhibitor of the Src/abl family | 20 |
| **Levetiracetam** | plays a role in the control of regulated secretion in neural and endocrine cells | 21 |
| **Tranylcypromine**[5] | belongs to a class of antidepressants monoamine oxidase inhibitors (MAOIs). | 22 |
| **HMN-214** | PLK inhibitor | 23 |
| **histamine** | immune responses, neurotransmitter | 24 |
| **dabrafenib** | chemotherapeutic, inhibitor of the associated enzyme B-Raf | 25 |

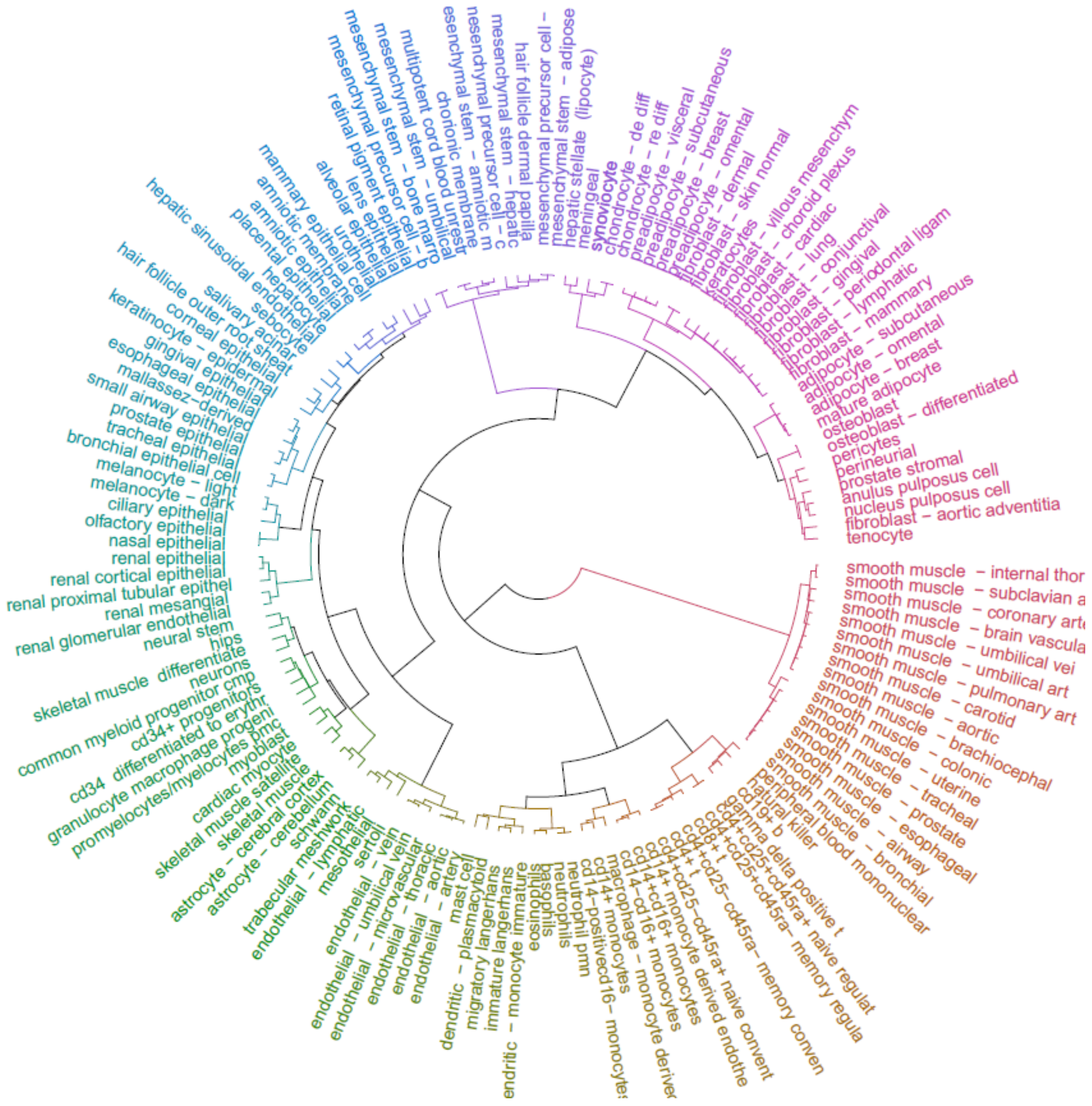

**Supplementary Figure 5:** Clustering of cell types based on the ontology distance. Affinity Propagation algorithm[8] was applied for the clustering.

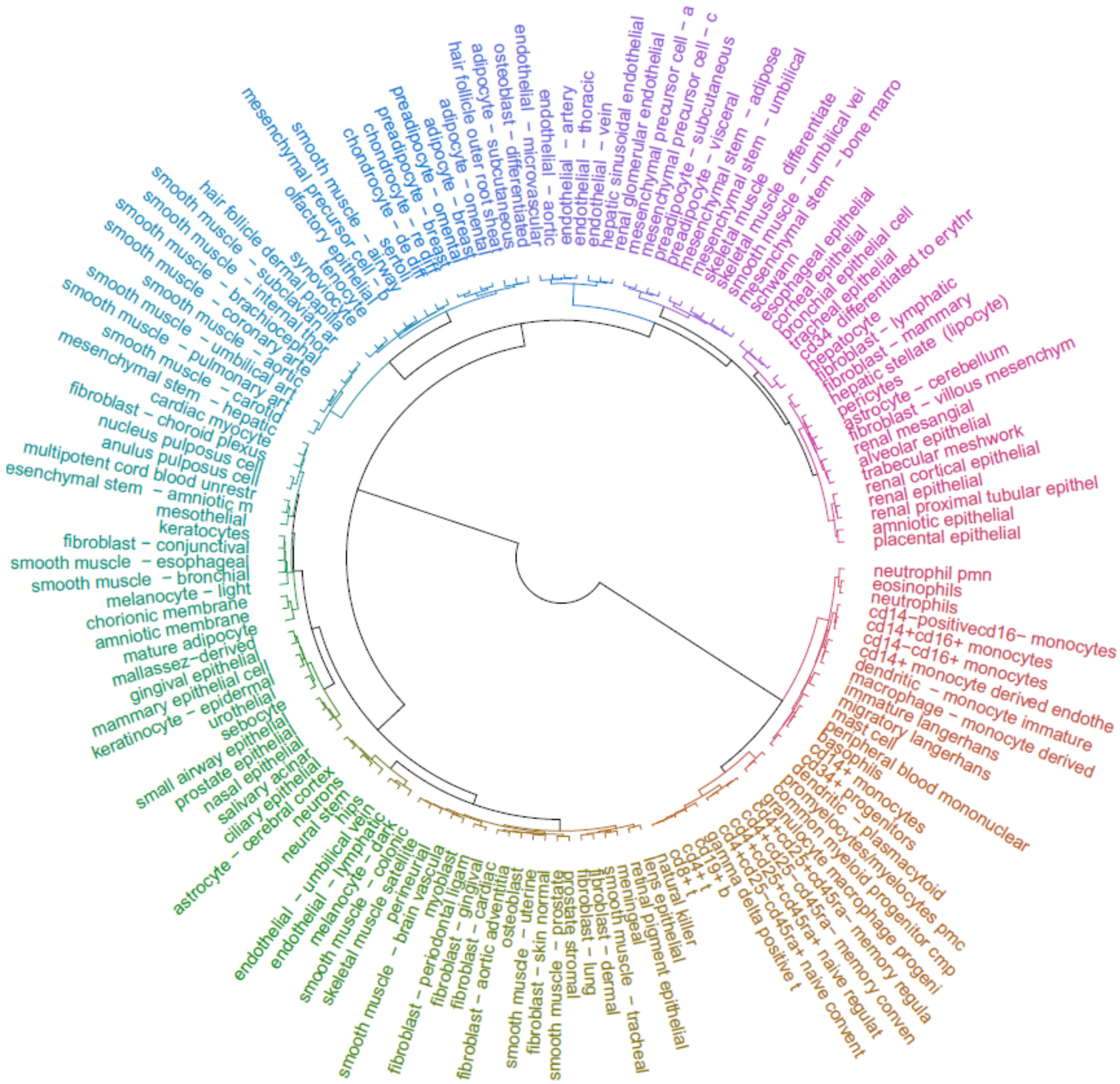


**Supplementary Figure 6:** Clustering of cell types based on the pathway distance. Affinity Propagation algorithm[8] was applied for the clustering.

**Supplementary Table 3:** The 69 meta-cell clusters created, based on both ontological and transcriptional similarity.

| Cluster | Included cell types |
|---|---|
| 1 | adipocyte - breast<br>adipocyte - omental<br>adipocyte - subcutaneous |
| 2 | alveolar epithelial cells<br>placental epithelial cells |
| 3 | amniotic epithelial cells |
| 4 | amniotic membrane cells<br>mammary epithelial cell<br>urothelial cells |
| 5 | anulus pulposus cell<br>nucleus pulposus cell |
| 6 | astrocyte - cerebellum |
| 7 | astrocyte - cerebral cortex |
| 8 | basophils<br>eosinophils<br>mast cell<br>neutrophil pmn<br>neutrophils |
| 9 | bronchial epithelial cell<br>esophageal epithelial cells<br>tracheal epithelial cells |
| 10 | cardiac myocyte |
| 11 | cd14+ monocyte derived endothelial progenitor cells<br>cd14+ monocytes<br>cd14+cd16+ monocytes<br>cd14+cd16- monocytes<br>cd14-cd16+ monocytes<br>peripheral blood mononuclear cells |
| 12 | cd19+ b cells<br>cd4+ t cells<br>cd4+cd25+cd45ra+ naive regulatory t cells<br>cd4+cd25+cd45ra- memory regulatory t cells<br>cd4+cd25-cd45ra+ naive conventional t cells<br>cd4+cd25-cd45ra- memory conventional t cells<br>cd8+ t cells<br>gamma delta positive t cells<br>natural killer cells |
| 13 | cd34 cells differentiated to erythrocyte lineage |
| 14 | cd34+ progenitors |
| 15 | chondrocyte - de diff |
| 1 | chondrocyte - re diff |
| 17 | chorionic membrane cells |
| 18 | ciliary epithelial cells<br>lens epithelial cells<br>retinal pigment epithelial cells |
| 19 | common myeloid progenitor cmp<br>granulocyte macrophage progenitor<br>promyelocytes/myelocytes pmc |

| | |
|---|---|
| 20 | corneal epithelial cells |
| 21 | dendritic cells - monocyte immature derived<br>immature langerhans cells<br>macrophage - monocyte derived<br>migratory langerhans cells |
| 22 | dendritic cells - plasmacytoid |
| 23 | endothelial cells - aortic<br>endothelial cells - artery<br>endothelial cells - microvascular<br>endothelial cells - thoracic<br>endothelial cells - vein<br>hepatic sinusoidal endothelial cells |
| 24 | endothelial cells - lymphatic<br>endothelial cells - umbilical vein |
| 25 | fibroblast - aortic adventitial<br>fibroblast - cardiac<br>fibroblast - gingival<br>fibroblast - periodontal ligament<br>fibroblast - skin normal |
| 26 | fibroblast - choroid plexus<br>fibroblast - villous mesenchymal<br>pericytes |
| 27 | fibroblast - conjunctival<br>fibroblast - dermal<br>fibroblast - lung<br>keratocytes |
| 28 | fibroblast - lymphatic<br>fibroblast - mammary |
| 29 | gingival epithelial cells<br>prostate epithelial cells<br>salivary acinar cells<br>small airway epithelial cells |
| 30 | hair follicle dermal papilla cells |
| 31 | hair follicle outer root sheath cells |
| 32 | hepatic stellate cells (lipocyte) |
| 33 | hepatocyte |
| 34 | hips<br>neural stem cells |
| 35 | keratinocyte - epidermal |
| 36 | mallassez-derived cells<br>sebocyte |
| 37 | mature adipocyte |
| 38 | melanocyte - dark |
| 39 | melanocyte - light |
| 40 | meningeal cells |
| 41 | mesenchymal precursor cell - adipose<br>mesenchymal precursor cell - bone marrow<br>mesenchymal precursor cell - cardiac |
| 42 | mesenchymal stem cells - adipose<br>mesenchymal stem cells - bone marrow |

| | |
|---|---|
| 43 | mesenchymal stem cells - amniotic membrane<br>multipotent cord blood unrestricted somatic stem cells |
| 44 | mesenchymal stem cells - hepatic |
| 45 | mesenchymal stem cells - umbilical |
| 46 | mesothelial cells |
| 47 | myoblast<br>skeletal muscle satellite cells |
| 48 | nasal epithelial cells |
| 49 | neurons |
| 50 | olfactory epithelial cells |
| 51 | osteoblast |
| 52 | osteoblast - differentiated |
| 53 | perineurial cells<br>prostate stromal cells |
| 54 | preadipocyte - breast<br>preadipocyte - omental<br>tenocyte |
| 55 | preadipocyte - subcutaneous<br>preadipocyte - visceral |
| 56 | renal cortical epithelial cells<br>renal epithelial cells<br>renal mesangial cells<br>renal proximal tubular epithelial cell |
| 57 | renal glomerular endothelial cells |
| 58 | schwann cells |
| 59 | sertoli cells |
| 60 | skeletal muscle cells |
| 61 | skeletal muscle cells differentiated into myotubes - multinucleated |
| 62 | smooth muscle cells - airway |
| 63 | smooth muscle cells - aortic<br>smooth muscle cells - brachiocephalic<br>smooth muscle cells - carotid<br>smooth muscle cells - coronary artery<br>smooth muscle cells - internal thoracic artery<br>smooth muscle cells - pulmonary artery<br>smooth muscle cells - subclavian artery<br>smooth muscle cells - umbilical artery |
| 64 | smooth muscle cells - brain vascular<br>smooth muscle cells - colonic<br>smooth muscle cells - esophageal<br>smooth muscle cells - tracheal |
| 65 | smooth muscle cells - bronchial |
| 66 | smooth muscle cells – prostate<br>smooth muscle cells - uterine |
| 67 | smooth muscle cells - umbilical vein |
| 68 | synoviocyte |
| 69 | trabecular meshwork cells |

**Supplementary Table 4:** Small molecules that were experimentally proved to facilitate various cell conversions and were predicted among the top 5% of the all drug profiles for the corresponding Meta-cells from the DECCODE single drug approach.

| Target Meta-cell | Small Molecule | Rank | Percentage |
|---|---|---|---|
| Astrocyte cells- Cerebral Cortex | Tranylcypromine[9] | 44 | 0.024887 |
| Hepatocyte cells | RG108[10] | 55 | 0.031109 |
| hIPS cells - Neural Stem cells | PD0325901[11, 12, 13] | 53 | 0.029977 |
| hIPS cells - Neural Stem cells | Tranylcypromine[14, 11] | 34 | 0.019231 |
| Mesenchymal Stem cells - Amniotic membrane - Multipotent Cord Blood Unrestricted Somatic Stem cells | PD0325901[15] | 59 | 0.033371 |
| Neurons | Y27632[16] | 7 | 0.003959 |
| Neurons | PD0325901[17] | 19 | 0.010747 |

**Supplementary Table 5:** Small molecules that were experimentally proved to facilitate various cell conversions and were predicted among the top 30 multidrug solutions for the corresponding Meta-cells from the DECCODE multi drug approach.

| Target Meta-cell | Small Molecule | Rank |
|---|---|---|
| Cardiac Myocyte cells | BIX01294[18] | 6 |
| hIPS cells - Neural Stem cells | Tranylcypromine[14, 11] | 18 |
| Neurons | PD032590[17] | 8 |
| Neurons | Y27632[16] | 19 |

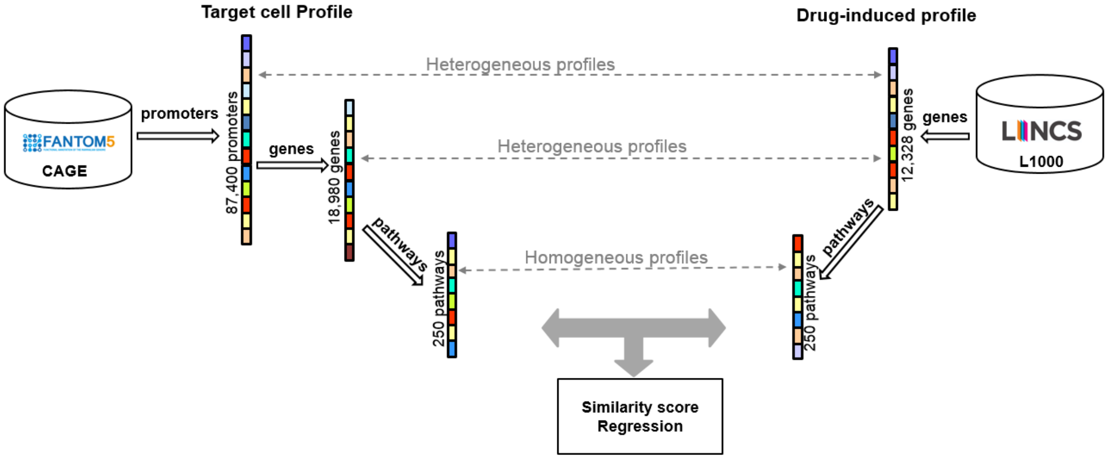


**Supplementary Figure 7:** Harmonization of expression profiles. Promoter-based target cell type profiles are converted to gene-based profiles. Gene-based profiles for both primary cells and drug treated cell lines are then converted to ~~pathway enrichment scores~~pathway-based expression profiles (PEPs) .

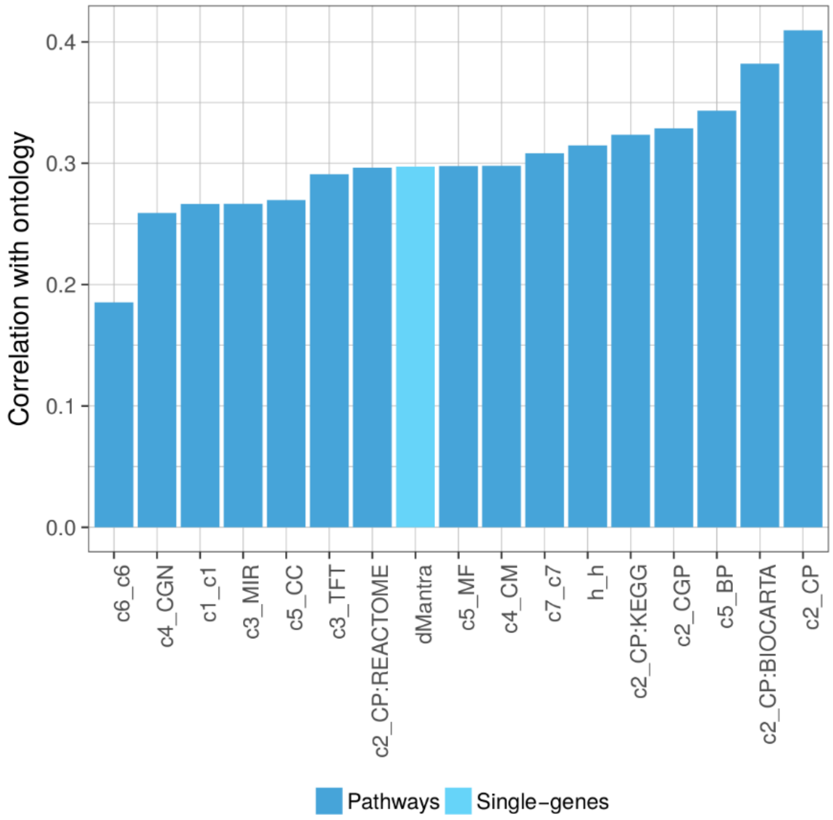


**Supplementary Figure 8:** Spearman correlation between pathway distances and ontology distance. Pairwise cell similarity obtained by the different pathway collections are evaluated against cell similarity obtained by the Cell Ontology annotation. Mantra distance[19] computed on single gene ranks is also tested against the ontology distance.

**Supplementary Table 6:** Pearson and Spearman correlation between fitted and observed PEPs in combinatorial treatment[20]. The multivariable linear regression model was applied in five different pathway collections. The values shown in the table are the mean performance after tenfold cross validation.

| | BP (4436 pathways) | | MF (901pathways) | | CC (580 pathways) | | TFT (615 pathways) | | C2_CP (250 pathways) | |
|---|---|---|---|---|---|---|---|---|---|---|
| | **Pearson** | **Spearman** | **Pearson** | **Spearman** | **Pearson** | **Spearman** | **Pearson** | **Spearman** | **Pearson** | **Spearman** |
| **Gefitinib-U0126** | 0.8038 | 0.7883 | 0.8685 | 0.8554 | 0.8401 | 0.8401 | 0.7929 | 0.8129 | 0.8538 | 0.8550 |
| **Gefitinib-Wortmannin** | 0.7460 | 0.7226 | 0.7538 | 0.7203 | 0.7947 | 0.6297 | 0.7276 | 0.7304 | 0.6814 | 0.6751 |
| **U0126-Wortmannin** | 0.6430 | 0.6211 | 0.6197 | 0.6095 | 0.6826 | 0.6502 | 0.6758 | 0.6336 | 0.6355 | 0.6080 |

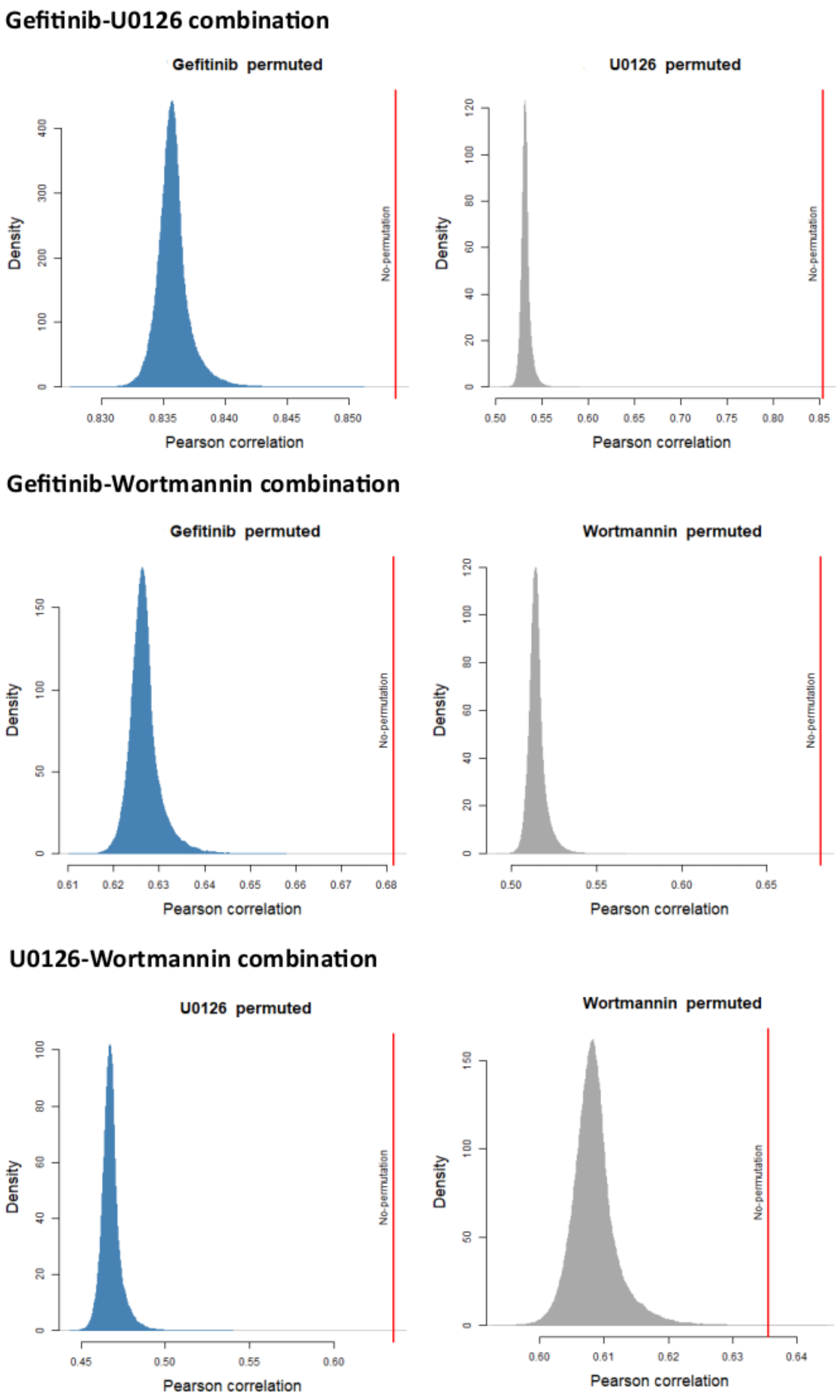


**Supplementary Figure 9:** Density plots of the Pearson correlation coefficients between observed and predicted values after the permutations of individual profiles for Gefitinib_U0126, Gefitinib_Wortmannin, and U0126_Wortmannin drug combinations using the C2_CP PEPs. The Pearson correlation coefficient achieved without permutation is also reported. Similar results were obtained for all the pathway collections of Supplementary Table 6.

### References

1. Napolitano, F., Sirci, F., Carrella, D. & di Bernardo, D. Drug-set enrichment analysis: a novel tool to investigate drug mode of action. *Bioinformatics* **32**, btv536 (2015).

2. Chen, G. *et al.* Blocking autocrine VEGF signaling by sunitinib, an anti-cancer drug, promotes embryonic stem cell self-renewal and somatic cell reprogramming. *Cell Res.* **24**, 1121–1136 (2014).

3. Yang, C. S., Lopez, C. G. & Rana, T. M. Discovery of nonsteroidal anti-inflammatory drug and anticancer drug enhancing reprogramming and induced pluripotent stem cell generation. *Stem Cells* **29**, 1528–1536 (2011).

4. Lin, T. & Wu, S. Reprogramming with Small Molecules instead of Exogenous Transcription Factors. *Stem Cells Int.* **2015**, 794632 (2015).

5. Di Stefano, B. *et al.* C/EBPα creates elite cells for iPSC reprogramming by upregulating Klf4 and increasing the levels of Lsd1 and Brd4. *Nat. Cell Biol.* **18**, 371–381 (2016).

6. Chang, M. Y. *et al.* Doxycycline enhances survival and self-renewal of human pluripotent stem cells. *Stem Cell Reports* **3**, 353–364 (2014).

7. Zhang, X., Simerly, C., Hartnett, C., Schatten, G. & Smithgall, T. E. Src-family tyrosine kinase activities are essential for differentiation of human embryonic stem cells. *Stem Cell Res.* 379–389 (2014). doi:10.1016/j.scr.2014.09.007

8. Frey, B. J. & Dueck, D. Clustering by Passing Messages Between Data Points. *Science (80-. ).* **315**, 972–976 (2007).

9. Tian, E. *et al.* Small-Molecule-Based Lineage Reprogramming Creates Functional Astrocytes. *Cell Rep.* **16**, (2016).

10. Zhu, S. *et al.* Mouse liver repopulation with hepatocytes generated from human fibroblasts. *Nature* **508**, 93–97 (2014).

11. Zhu, S. *et al.* Reprogramming of Human Primary Somatic Cells by OCT4 and Chemical Compounds. *Cell Stem Cell* **7**, 651–655 (2010).

12. Wang, Q. *et al.* Lithium, an anti-psychotic drug, greatly enhances the generation of induced pluripotent stem cells. *Cell Res.* **21**, 1424–1435 (2011).

13. Lin, T. *et al.* A chemical platform for improved induction of human iPSCs. *Nat. Methods* **6**, 805–808 (2009).

14. Li, W. *et al.* Generation of Human Induced Pluripotent Stem Cells in the Absence of Exogenous *Sox2*. *Stem Cells* **27**, N/A-N/A (2009).

15. Lai, P.-L. *et al.* Efficient Generation of Chemically Induced Mesenchymal Stem Cells from Human Dermal Fibroblasts. *Sci. Rep.* **7**, 44534 (2017).

16. Hu, W. *et al.* Direct Conversion of Normal and Alzheimer's Disease Human Fibroblasts into Neuronal Cells by Small Molecules. *Cell Stem Cell* **17**, (2015).

17. Dai, P., Harada, Y. & Takamatsu, T. Highly efficient direct conversion of human fibroblasts to neuronal cells by chemical compounds. *J. Clin. Biochem. Nutr.* **56**, 166–170 (2015).

18. Cao, N. *et al.* Conversion of human fibroblasts into functional cardiomyocytes by small molecules. *Science* **1502**, (2016).

19. Iorio, F. *et al.* Discovery of drug mode of action and drug repositioning from transcriptional responses. *Proc. Natl. Acad. Sci.* **107**, 14621–14626 (2010).

20. Rapakoulia, T. *et al.* Genome-scale regression analysis reveals a linear relationship for promoters and enhancers after combinatorial drug treatment. *Bioinformatics* **33**, (2017).
